# Evaluating metabarcoding to analyse diet composition of species foraging in anthropogenic landscapes using Ion Torrent and Illumina sequencing

**DOI:** 10.1101/295758

**Authors:** Marie-Amélie Forin-Wiart, Marie-Lazarine Poulle, Sylvain Piry, Jean-François Cosson, Claire Larose, Maxime Galan

## Abstract

DNA metabarcoding of faecal samples is being successfully used to study the foraging niche of species. We assessed the ability of two benchtop high-throughput sequencing (HTS) platforms, to identify a large taxonomic array of food items from domestic cats *Felis silvestris catus*, including prey and human-related food taxa (pet food and leftovers leaving undetectable solid remains in faeces). Scats from a captive feeding trial (*n*=41) and from free-ranging individuals (*n*=326) were collected and analysed using a cyt*b* mini-barcode in independent PCR duplicates on the Ion PGM and the MiSeq platforms. Outputs from MiSeq were more sensitive and reproducible than those from Ion PGM due to a higher sequencing depth and sequence quality on MiSeq. DNA from intact prey taxa was detected more often (82% of the expected occurrences) than DNA from pet food (54%) and raw fish and meat (31%). We assumed that this variability was linked to different degree of DNA degradation: The Ion PGM detected significantly less human-linked food, birds, field voles, murids and shrews in the field-collected samples than the MiSeq platform. Pooling the replicates from both platforms and filtering the data allowed identification of at least one food item in 87.4% of the field-collected samples. Our DNA metabarcoding approach identified 29 prey taxa, of which 25 to species level (90% of items) including 9 rodents, 3 insectivores, 12 birds and 1 reptile and 33 human-related food taxa of which 23 were identified to genus level (75% of items). Our results demonstrate that using HTS platforms such as MiSeq, which provide reads of sufficiently high quantity and quality, with sufficient numbers of technical replicates, is a robust and non-invasive approach for further dietary studies on animals foraging on a wide range of food items in anthropogenic landscapes.

## Introduction

With increasing urbanisation and loss of natural habitat, many species have adapted to human-dominated landscapes by shifting their dietary niche to include human-processed food items^1^. For instance, scavengers such as gulls *Larus spp.*, ravens *Corvus spp.*, Eurasian badgers *Meles meles*, opossums *Didelphis virginiana*, racoons *Procyon lotor*, or wild boar *Sus scrofa*, as well as opportunistic generalist carnivores such as coyotes *Canis latrans*^2^, or foxes *Vulpes spp.*^3,4^, presently have dietary compositions that differ substantially from their natural diet. Several large carnivores such as bears, wolves and hyenas have also altered their diet to include a significant proportion of human-processed products (see Bateman and colleagues^5^ for a review). Understanding to what extent species in close contact with humans may have changed their diet to include human-processed items is critical both from a conservation perspective to determine the likelihood of human-wildlife conflicts, and from an evolutionary perspective to determine species plasticity in foraging niches in a context of global change. Thus, accurately identifying the large diversity of food items in the diet composition of studied species, including those from human-linked food sources, is necessary for ecologists.

Foraging ecology studies generally rely on the preservation of hard remains found in the faeces, pellets, or stomach contents of studied species to determine their diet, using visual recognition of morphologically features by macro- or microscopic methods^6^. Whereas useful, these techniques generally result in poor resolution in terms of taxa identification or preclude the identification of food items that do not leave behind hard remains such as pet food or leftovers only composed by soft tissues^7,8^. In contrast, DNA metabarcoding has been demonstrated as an accurate method for diet analysis, using faecal samples as a source of DNA^9^. As a result, DNA metabarcoding using high-throughput sequencing (HTS) has become a valuable tool for ecological research^10^. This approach uses PCR amplification of a genetic barcoding marker to analyse mixtures of DNA isolated from biological samples, for example enabling detection of predator and prey DNA from previously unidentified samples.

Metabarcoding in HTS technologies allows DNA to be read from multiple templates for hundreds of samples in the same sequencing run. A subunit of a mitochondrial gene – for example, a 136-bp mini-barcode on the cytochrome *b* gene (cyt*b*) – is enough to distinguish between closely related rodent species even in degraded or ancient DNA samples^11^. By comparing obtained sequences to a standard reference library of taxonomically identified organisms (e.g. Bold or GenBank), the taxa or species present in a sample can be identified with high confidence. In this way, metabarcoding of faecal samples allows ingested food items to be identified by characterizing the barcode sequences in the samples^12^. This technique has been successfully used to analyse the diet of species of birds^13^, mammals^11,14,15^, reptiles^16,17^ and fish^18^.

However promising, metabarcoding for dietary analyses is not completely free of limitations or bias (see Table 1 in Galan and colleagues^19^ for a review). Contamination from alien DNA during fieldwork or cross-contamination during lab procedures can occur, resulting in false positive results^20^. Furthermore, the library preparation protocol, the targeted DNA region, the primer choice and the sequencing platform can all affect the identification of the composition of environmental samples^21^. The degree of DNA degradation in an analysed sample also influences the sensitivity of PCR-based methods^22^. In addition, the sequencing of PCR products from samples that contain predator DNA could reduce the potential detection of the prey sequences^15^. As a result of these potential biases, alone or in combination, the relative abundance of some species in faecal samples can be under- or overestimated in a DNA analysis, while other species may not be amplified and so remain absent from the bioinformatics output^23^. Although several studies have described most of these challenges, to our knowledge the specific use of HTS platforms to analyse samples from a dietary perspective has never been evaluated.

**Table 1:**
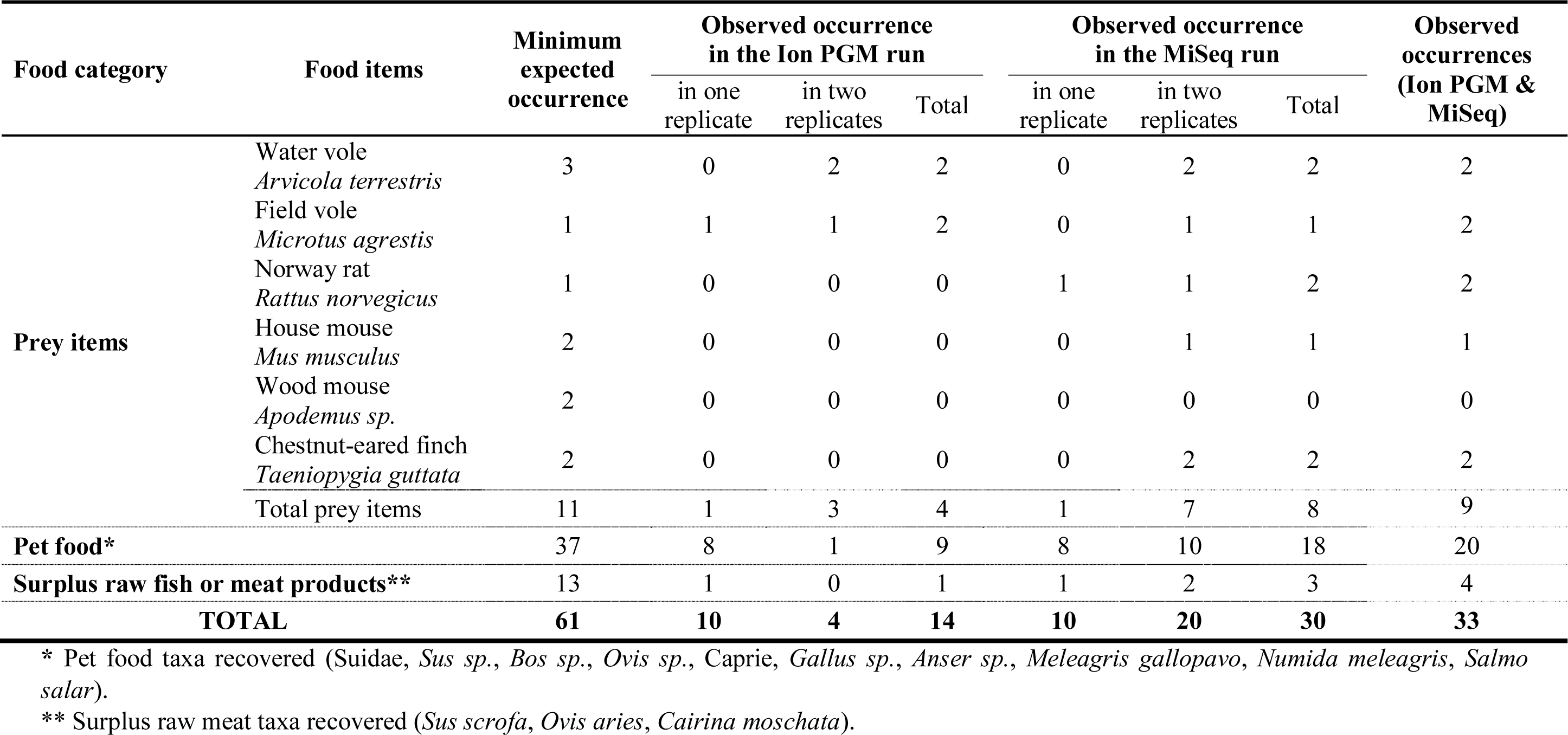
Outputs from Ion PGM and MiSeq platforms during the calibration study. The minimum expected occurrences and the actual observed occurrences of food items in HTS runs of 41 faecal samples collected from a housebound cat. The observed occurrences are shown according to the HTS platform used, the PCR replicates and the pooled occurrences from both platforms.

The main objective of this work was to assess the ability of benchtop HTS platforms in identifying a large taxonomic array from the diet of species that potentially contain human-processed products. To maximise the variability in dietary DNA samples obtained and ensure the presence of human-linked food items, we focused on a highly generalist carnivore species, the domestic cat *Felis silvestris catus*. Cats are also a particularly interesting model to test for metabarcoding sensitivity as in addition to human-processed products, free-ranging individuals are known to feed on a wide diversity of prey items including rodent species, birds, reptiles, amphibians and insects^24,25^. Specifically in this study we investigated the sensitivity of the two most popular sequencers available for metabarcoding analyses, the Life Technologies Ion PGM (Personal Genome Machine) platform and the Illumina MiSeq platform, in identifying food items from faecal DNA having undergone different degrees of degradation (fresh prey items versus processed food versus surplus raw fish or meat products). This was investigated for (1) ecological reasons, i.e. understand the relative contributions of wildlife versus food scraps provided by or scavenged from humans to the diet, and so illuminate predator ecology and (2) technical reasons, i.e. understand the composition of food items which affects the detectability of DNA. This was achieved by collecting faecal samples from (1) a cat fed on a mixed diet of a known composition in an initial calibration study, and (2) free-ranging individuals sampled from a rural area. Since all samples were analysed in independent PCR duplicates on each HTS platform, this allowed us to compare the reproducibility of diet results both within the platform and among platforms.

## Results

### Sequencing results

The Ion PGM and MiSeq platforms provided 4,042,017 and 4,042,606 raw sequences passing the filter respectively. For the Ion PGM reads, the percentage of bases with a quality score of 30 (Q30: incorrect base call probability of 1 in 1000) or higher was 61.6%; for the MiSeq reads, the percentage of bases with a quality score of 30 or higher was 96.9%. For the Ion PGM run, a high proportion of truncated sequences without reverse tags was observed (2,490,973 reads < 126bp excluding tags and primers) due to a significant decrease in quality at the end of the sequences. After demultiplexing, 688,530 (using Ion PGM) and 2,509,592 (using MiSeq) sequences of cyt*b* were assigned to the faecal samples and were identified to the closest taxonomic rank (genus or species) using two out of three sliding similarity thresholds (i.e. 100% or 98% for prey taxa and 98% or 95% for food processed taxa).

In the calibration study (*n* = 41 faecal samples), we obtained a high proportion of reads from the host (i.e., cat haplotypes): 41,129 reads (87.5 %) for Ion PGM and 235,673 reads (98.5%) for MiSeq at the 95% similarity threshold. There were also a high proportion of unidentified sequences: 5,069 reads (10.8 %) for Ion PGM and 1,434 reads (0.6%) for MiSeq at the 95% similarity threshold. Accordingly, informative reads for the diet accounted for 1.6% for Ion PGM (756 reads) and 0.9% (2,147 reads) for MiSeq at the 95% similarity threshold. In the study of field-collected data (*n* = 359 faecal samples), sequences from predators (cat, but also Canids and Mustelids, see below) represented 60% of the total read count using Ion PGM, and 88% of the total read count using MiSeq. For both platforms, a high proportion of reads from the host (301,329 reads [51%] for Ion PGM and 1,690,359 reads [83%] for MiSeq) and unidentified sequences (70,050 reads [12%] for Ion PGM and 125,086 reads [6%] for MiSeq, with a ≤ 95% identity using GenBank sequences) were obtained. Informative reads for the diet accounted for 28% for Ion PGM (165,261 reads) and 6% (116,423 reads) for MiSeq at the 95% similarity threshold. As found in a previous study by Deagle and colleagues^26^, some human DNA sequences were also present, accounting for 0.06% and 0.6% of the sequences generated by Ion PGM and MiSeq respectively.

### DNA metabarcoding on known mixed-diet samples

We estimated the false assignment rate (R_FA_) in our experiment by looking for reads from food taxa absent from calibration samples but present in the field-collected samples. The occurrence of reads belonging to this variant in these samples constituted a miss-tagging event. In the MiSeq run, one out of 1934 sequences of a *Myodes glareolus* haplotype (n = 1, resulting in a rate of 0.0517) was detected during the last 21 days of the calibration study despite the fact the housebound cat was fed only with pet food and raw fish or meat (see Table S1 and Table S5 for details). Accordingly, the false-assignment rate (R_FA_) in a sequencing run was set at 0.05%. Negative extraction and PCR controls contained reads belonging to a human haplotype, non-assigned ones and a cat haplotype but none were from food taxa. Once data was filtered using T_FA_ and replicates, reads were no longer consistent and informative.

Of the 11 distinct prey items fed to the housebound cat, 8 were detected using MiSeq (73%), and 7 of these were detected in both of the PCR replicates from the relevant sample (Table 1 and Table S1). In contrast, only 4 expected prey items were detected using Ion PGM (36%), of which 3 were detected in both PCR replicates from the relevant sample. MiSeq was the only platform to detect the Norway rat, the house mouse and the Chestnut-eared finch (Table 1). Of the 37 expected pet food occurrences, 18 were detected using MiSeq (49%), and 10 of these were detected in both of the PCR replicates from the relevant sample (Table 1 and Table S1). However, only 9 pet food occurrences were detected using Ion PGM (24%), of which only 1 was detected in both PCR replicates from the relevant sample. Of the 13 expected raw fish and meat occurrences, 3 were detected using MiSeq (23%), and 2 of these were detected in both of the PCR replicates from the relevant sample (Table 1 and Table S1). In contrast, only 1 not replicated raw meat occurrence was detected using Ion PGM (8%).

Both platforms were better at detecting DNA from prey items (82% of the expected occurrences) than DNA from pet food (54%), or from raw fish and meat (31%) (Table 1). Independent of the food item considered, the percentage of faecal samples in which the expected DNA of a food item was detected (see Table S1) was higher for the MiSeq run (25/27, 93%) than for the Ion PGM run (14/27, 52%) (Table 1). Combining the results of the HTS runs from both platforms, the sequencing of food item DNA, in the faeces collected from the housebound cat, succeeded in 66% of the samples (27 out of 41 samples).

Of the six expected prey species, the wood mouse *Apodemus sp.* was not detected, while at least one of the two HTS platforms sequenced four other rodent species (water vole, field vole, Norway rat and house mouse) and a bird species (Table 1). DNA from the Norway rat and field vole, each expected in one scat, as with the other food items, was recovered in two consecutive faeces. DNA from the house mouse was poorly detected (i.e. only four informative reads from the MiSeq run), and the wood mouse was not detected at all. A delay of 12 to 24 hours between prey ingestion and the occurrence of their remains in the scat was identified for each prey species eaten and detected by both methods.

The proportion of sequences (Ion PGM + MiSeq) associated with the different food item categories did not match the proportion of the food item consumed (p < 0.001; Fig. 1a), even when the platforms were considered independently (Fig. 1b and 1c). Prey items were considerably overrepresented in comparison to other diet items: they accounted for 66% of the sequence reads assigned to food items, whereas they represented only 5% of the biomass ingested by the cat. Within the prey item category, rodent and bird biomass (90.4% and 9.6% respectively) were representative of their read proportions (95.7% and 4.3% respectively, p > 0.05). Conversely, human-linked food items (pet food and raw fish or meat) were considerably underrepresented: pet food accounted for 33% and raw fish or meat 0.5% of the sequence reads, whereas these respectively represented 83% and 12% of the biomass ingested.

**Figure 1:**
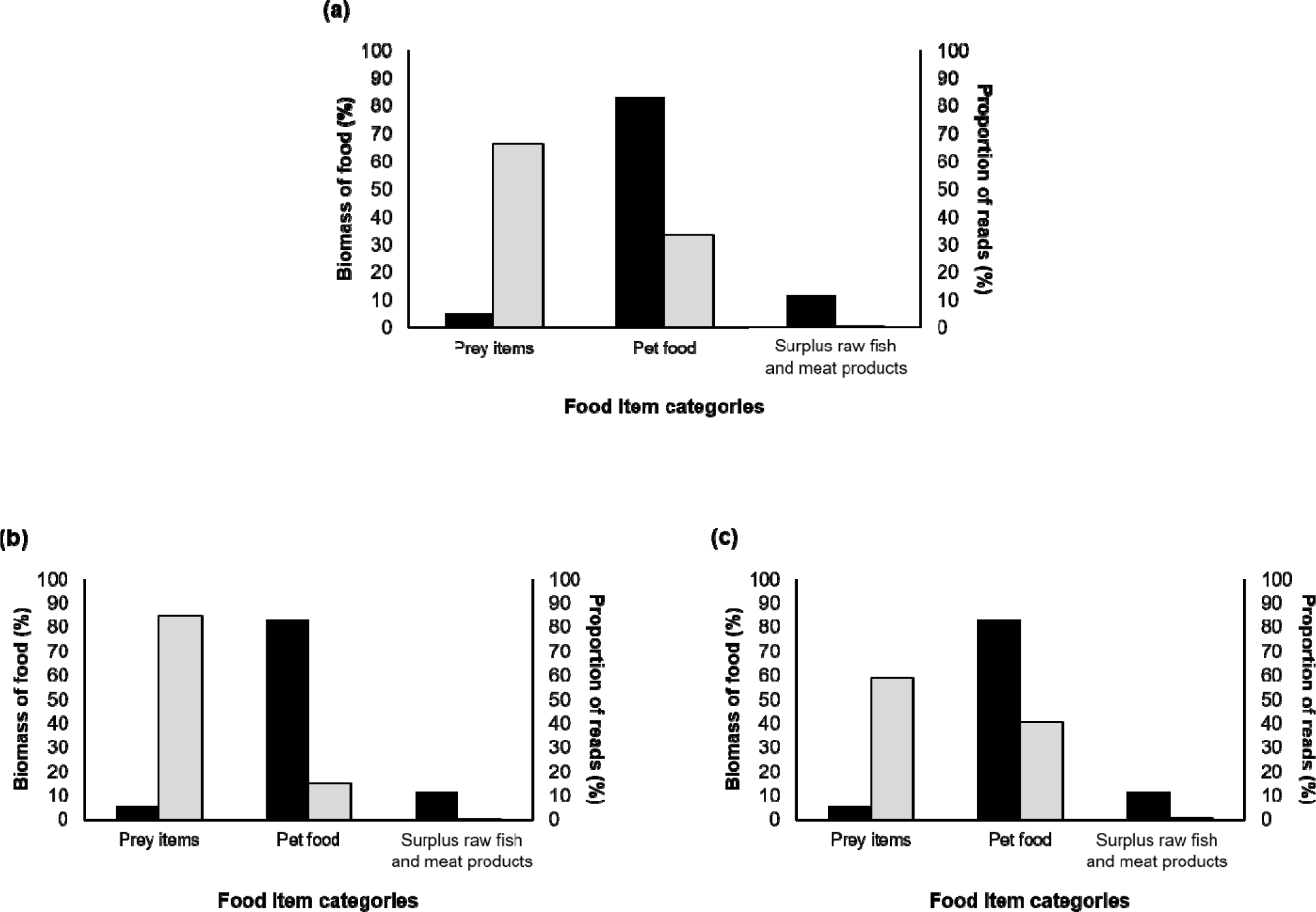
The relationship between detection rate and variability in DNA degradation. Comparison between the proportion of biomass of the three food categories fed to the housebound cat (black) versus the overall proportion of Sequence reads assigned to that food category (light grey) from 41 scat samples for (a) pooled replicates (Ion PGM + MiSeq runs), (b) Ion PGM replicates only, and (c) MiSeq replicates only.

### DNA metabarcoding on field-collected samples

Comparing predator identification using cat-specific PCR amplification and DNA metabarcoding, 2.4% of faecal samples (10/415) identified as cat faeces using cat-specific PCR amplification, were found to be from another predator species using DNA metabarcoding; 5% of faecal samples (20/415) not identified as cat faeces using cat-specific PCR amplification were found to be from a cat using DNA metabarcoding. Of the 359 putative felid faecal samples collected, 90.8% (326/359) were confirmed as originating from domestic cats using sequencing. The others were determined to be the scats of Canids (4.5% from the domestic dog *Canis lupus familiaris* and 3.9% from the red fox,) or Mustelids (0.6% from the Eurasian badger *Meles meles* and 0.3% from *Martes sp.*). All a priori DNA metabarcoding identifications of predators were confirmed unambiguously by both HTS platforms and by the four PCR replicates.

Of the 180 observed occurrences of prey items detected with MiSeq, 72% were identified in both of the PCR replicates compared to 65% for Ion PGM (103/159). Of the 189 observed occurrences of human-linked food items (pet food and scraps) detected with MiSeq, 48% were identified in both of the PCR replicates compared to 39% for Ion PGM (103/159). The two platforms jointly confirmed 72.2% (143/198) of the occurrences of prey items, and 61% (115/189) of the occurrences of human-linked food. In terms of the cat faecal samples, Ion PGM detected at least one food item in 73.6% (240/326) of the investigated samples, and MiSeq detected at least one food item in 83.1% (271/326) of cases. Combining the results from both HTS platforms, 87.4% (285/326) of the investigated samples contained at least one identified food item.

A likelihood ratio test applied to the model selected for food item detection indicated that the model without random effects was significantly better than the one with random effects (L = 870.09, df = 1, p > 0.05). The top-ranked model included fPlatform and fFoodItem and the interaction between these two variables (R² = 0.17, Table S4). Ion PGM detected significantly less human-linked food (−2.01 ± 0.31 SE on the log odds ratio scale, p < 0.001), birds (−4.98 ± 1.10 SE, p < 0.001), field voles (−1.50 ± 0.72 SE, p < 0.05), murids (−3.30 ± 1.10 SE, p < 0.01) and shrews (−2.60 ± 1.24 SE, p < 0.05) than MiSeq.

A total of 774 different occurrences of food items were identified and classified as prey items or human-linked food (Table 2 and Table S2). A total of 29 different prey taxa were identified, of which 25, representing 90% of the prey items (287/319), were identified up to the species level (9 rodents, 3 insectivores, 12 birds and 1 reptile) while the remaining 10% were identified to the genus level (32/319, Table 2 and Table S2). One reptile item was identified as being the slow worm *Anguis fragilis*. A total of 455 occurrences that were related to human-linked food items were detected (Table 2 and Table S2). Game species were also detected in addition to the expected occurrences of pet food, food leftovers and hunting remains. A total of 33 different taxa related to human-linked food were identified, of which 17 were identified without ambiguity up to the species level (36%, 165/455) or 23 up to the genus level (75%, 342/455), the remaining 25% of human-related food items were identified to higher taxonomic ranks (family, order or superorder; 113/455, Table 2 and Table S2).

**Table 2:**
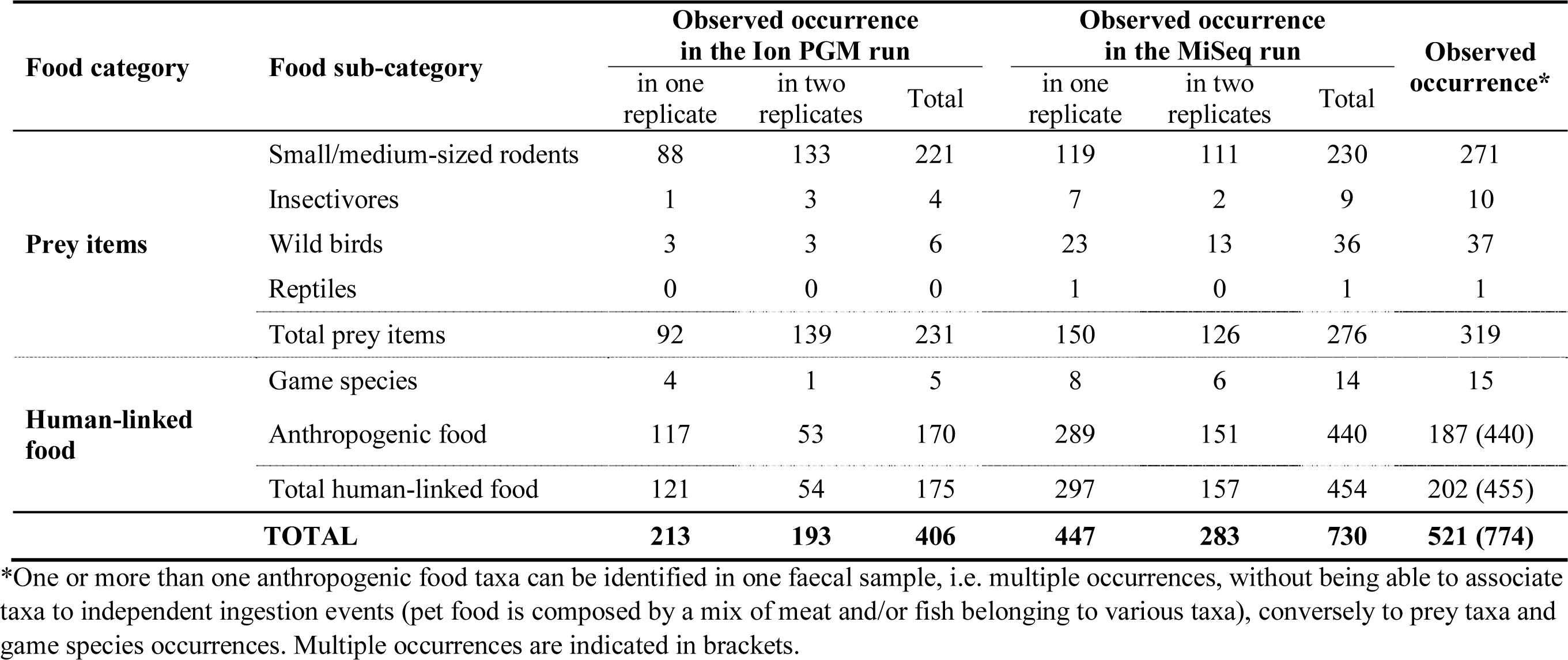
Outputs from Ion PGM and MiSeq platforms during the field study. Observed occurrences of food items in 326 field-collected faecal samples from free-ranging cats. The observed occurrences are shown according to the HTS platform used and the PCR replicates. The final observed occurrence corresponds to the pooled results from the Ion PGM and MiSeq runs. See Table S2 for more details about food sub-categories.

| Food category | Food sub-category | Observed occurrence<br>in the Ion PGM run |  |  | Observed occurrence<br>in the MiSeq run |  |  | Observed<br>occurrence* |
| --- | --- | --- | --- | --- | --- | --- | --- | --- |
|  |  | in one<br>replicate | in two<br>replicates | Total | in one<br>replicate | in two<br>replicates | Total |  |
| **Prey items** | Small/medium-sized rodents | 88 | 133 | 221 | 119 | 111 | 230 | 271 |
|  | Insectivores | 1 | 3 | 4 | 7 | 2 | 9 | 10 |
|  | Wild birds | 3 | 3 | 6 | 23 | 13 | 36 | 37 |
|  | Reptiles | 0 | 0 | 0 | 1 | 0 | 1 | 1 |
|  | Total prey items | 92 | 139 | 231 | 150 | 126 | 276 | 319 |
| **Human-linked**<br>**food** | Game species | 4 | 1 | 5 | 8 | 6 | 14 | 15 |
|  | Anthropogenic food | 117 | 53 | 170 | 289 | 151 | 440 | 187 (440) |
|  | Total human-linked food | 121 | 54 | 175 | 297 | 157 | 454 | 202 (455) |
| **TOTAL** |  | **213** | **193** | **406** | **447** | **283** | **730** | **521 (774)** |

\*One or more than one anthropogenic food taxa can be identified in one faecal sample, i.e. multiple occurrences, without being able to associate taxa to independent ingestion events (pet food is composed by a mix of meat and/or fish belonging to various taxa), conversely to prey taxa and game species occurrences. Multiple occurrences are indicated in brackets.

## Discussion

The use of the short cyt*b* mini-barcode provided a good sensitivity of food item detection, particularly for prey items. Galal-Khallaf and colleagues^18^ highlighted the advantage of amplifying a short mini-barcode when employing a DNA metabarcoding approach to degraded DNA in highly processed food, based on 454 Next Generation Sequencing (NGS), to detect fish species in commercial fish feed samples. However, in our study non-reproducible occurrences between replicates were observed using Ion PGM and MiSeq runs independently (86% for Ion PGM and 56% for MiSeq runs during the calibration study, and 48% for Ion PGM and 40% for MiSeq runs during the field study). Using a non-arbitrary threshold (T_FA_) that is calculated according to variant abundance for each sequencing run allows the mitigation of the occurrence of false positives^27^ while leading occasionally to false negative results (e.g. 0 and 5 reads of *Bos sp.* at the 95% similarity threshold for Ion PGM replicates [T_FA*Bos*_= 5 reads] and no reads of *Bos sp.* at the 95% similarity threshold for MiSeq replicates [T_FA*Bos*_= 17 reads], respectively during the calibration study). Such findings emphasize the need to generate sufficient numbers of technical replicates (at least four were required in the present study) and to increase the sequencing depth to detect rare or degraded DNA, in order to limit the presence of false negatives as well as to alleviate inconsistent results likely to be false positives^28^.

Results from the calibration study revealed that DNA related to processed food is rarely recovered in duplicates and is less likely to be detected than the DNA of prey items by both HTS platforms. Given that there is no critical 3’-terminal mismatches between PCR primers and DNA targets (i.e., underlying a bias of detection for particular taxa), this result might be linked to a DNA integrity gradient between prey items, pet food and raw fish and meat. Prey items were frozen immediately after death during the calibration study, and in the field study they were probably the result of hunting rather than scavenging behaviour^25^, ensuring maximum DNA integrity before ingestion. In addition, in this food category, hard parts (e.g. bones versus flesh or other soft parts) limit DNA degradation during digestion, and so partly solve problems of variable digestibility^7^. In contrast, the stages of food processing (e.g. cooking, dehydration, heat sterilization, etc.) involved in producing pet food induce a higher rate of DNA degradation than for prey items, leading to damaged and degraded DNA^22,29^. However, food processing also suppresses the activity of micro-organisms such as bacteria, so the integrity of the DNA remains the same regardless of the duration of food storage before ingestion. In the case of the category of raw fish and meat, which is not processed, DNA integrity depends on the length of storage and the activity of micro-organisms. In this study, the use of surplus raw fish and meat products stored in a supermarket refrigerator for several days/weeks might explain the low success of DNA detection related to this food category. The link between food category and DNA integrity could be confirmed through a calibration study including the evaluation of DNA integrity and a DNA metabarcoding experiment on various food item categories before and after ingestion. Subsequently, analysing the rate of detection for a given food item category before ingestion could be used to reveal problems of PCR amplification bias (e.g. due to primer mismatches or PCR inhibitors in the food). Furthermore, we could not measure the variability of DNA detection of specific food items from one cat to another, since we performed this experiment only with one individual. This variability could be evaluated through an additional calibration study including several cats fed with the same diet. Such a protocol would allow the exploration of DNA metabarcoding limits, by considering the variability of DNA detection, regarding the DNA integrity of food items, linked to an individuals’ digestion capacities (e.g. usual diet, age, sex, health status and even breed of cat).

In addition to variability in the DNA detection of food items, variations in the ability of the different HTS platforms to detect food items were identified. Overall, during the calibration study, MiSeq was more effective at detecting expected food items and duplicating this (93% of expected items were detected, of which 71% were duplicate-confirmed) than Ion PGM (52% of expected items were detected, of which 29% were duplicate-confirmed). A similar pattern was observed in the analysis of the field-collected faecal samples. Furthermore, MiSeq was also able to detect more human-linked food, as well as species of bird, field vole, murids and shrews than Ion PGM in field collected-samples. This outcome could be most easily explained by the difference in the number of sequence reads assigned to the samples between the two HTS platforms (i.e. low sequencing depth for the Ion PGM platform), linked with the better sequencing quality of the MiSeq platform.

Another strong sequencing bias was noticed in results from the Ion PGM platform, particularly for species haplotypes that included short homopolymer repeats. In our case, a loss of quality at the end of the cyt*b* mini-barcode, and consequently to the presence of several short homopolymer stretches in tandem (e.g. the last 22 bases of the mini-barcode from *Felis silvestris catus*: 5’-CCC TTT AAA TAC CCC TCC CCA T-3’) explained a high proportion of truncated sequences without reverse tags. This bias has been previously observed in other public datasets generated by this platform and suggests that it is a pervasive problem^30^. This may also partly explain the bias found between biomass and number of reads, leading to a lower representation of homopolymer-rich haplotype sequences in the DNA metabarcoding studies using Ion Torrent technology (e.g. Deagle and colleagues^31^).

The proportion of reads from prey items or human-linked food was not related to the ingested biomass of these food categories in the housebound cat diet. On a mass-to-mass basis, prey item DNA was better detected than DNA associated with human-linked food. This result was not related to a taxonomic bias introduced by primer-template mismatches, but rather related to food item properties. This discrepancy is likely to be associated with the variable digestibility of different food items and/or DNA quality (see above). In line with previous studies^31-33^, these results reinforce the preference for a qualitative approach that involves creating a list of food items consumed and their frequency of occurrence^12,26^ rather than relating relative number of reads to ingested biomass^34^, particularly in studies on species with mixed diets (i.e. preys and processed food taxa). This issue has been highlighted in several recent DNA-based dietary studies investigating the COI barcode region^35^ or a more conserved mitochondrial region such as the mitochondrial 16S rRNA gene^36^. Deagle and colleagues^31^ also noted a lack of relationship between biomass and the proportion of reads when comparing the proportion of three fish species detected in seal scat to the proportion of these species in a constant diet. One solution would be to work with different metrics according to the observed food items comprised in the diet (e.g. occurrence data for preys and processed food taxa comparison versus relative read abundance for within preys taxa comparison)^37^. An alternative would be to use conserved markers such as those targeting the mitochondrial 12S rRNA or the nuclear 18S rRNA^38^, but these provide limited taxonomic resolution for vertebrates compared to markers targeting mitochondrial cytochrome *b*. One way to increase pet food detection in dietary studies would be to target not only vertebrate and invertebrate components of the diet but also cereals contained in pet food. DNA metabarcoding that targets different genes, including those of plants, has been successfully used on an omnivorous carnivore, the brown bear *Ursus arctos*^14^. DNA identified in faeces could also reflect secondary predation cases, e.g. insects consumed by shrews or seeds ingested by rodents^11,39^.

Another bias related to an approach based on amplifying a cyt*b* 136-bp mini-barcode without a blocking oligonucleotide on the targeted predator species is that it can lead to an over-representation of the sequences assigned to the predator in the PCR product. As a result, some samples can exclusively yield predator sequences by inhibiting the amplification of rare DNA reads (e.g. prey items). This phenomenon is probably exacerbated because predator DNA has higher integrity than DNA originating from human-linked food. This hypothesis is consistent with the high proportion of cat haplotypes in the present study (87.5% for Ion PGM and 98.5% for MiSeq when faecal samples were rapidly stored, and 51% for Ion PGM and 83% for MiSeq in more hazardous sample conservation). Of the field-collected faecal samples, 18% produced only cat sequences. Similarly, in a study by Shehzad and colleagues^15^, 21% of faecal samples were found to be entirely composed of leopard cat *Prionailurus bengalensis* DNA when investigating leopard cat diet using metabarcoding without a blocking oligonucleotide on the targeted predator species. The high proportion of predator sequences in the PCR product reduces the sensitivity of HTS platforms. This phenomenon contributes to explaining the relatively low sequencing success rate obtained with Ion PGM, for which the total number of reads was lower than with MiSeq. However, increasing sequencing depth (i.e. increasing the number of informative reads) is not the only determining factor to consider when improving taxa detection. Producing technical replicates is an effective solution to overcome stochastic DNA amplification from certain taxa, thus improving the overall diet as a whole (see Alberdi and colleagues^40^ and the present study).

In line with D’Amore and colleagues^21^, these results indicate that studies using DNA metabarcoding for conservation management purposes should evaluate their outputs in comparison with a calibration study or at least with biological expectations. It should be noted that our calibration study also indicated underestimation (e.g. small murids) and overestimation (e.g. DNA from large species [Norway rat] or numerously ingested prey species [field vole] found in more than one sample) of food items in cat diets; this could be linked to the number of food taxa in samples^37^ or targeted species biology (i.e. pyloric retention in Felids). Several previous studies also support this pattern^41-44^.

This study also confirmed that species assignment is subject to errors related to sequences incorrectly attributed to the wrong species in GenBank (e.g. closely related rodents/birds). A precise and full knowledge of the geographical ranges of species can even partially correct such errors. DNA metabarcoding is mainly used to investigate faeces with no previous knowledge about the diet of the targeted species; a reference database containing voucher specimens from the study area would be useful to validate the detection of prey species. One alternative would be to use tools such as PrimerMiner^45^ to obtain relevant sequence data for targeted primer thus reducing biases introduced by the different number of available sequences per species and evaluate primers *in silico*.

When dealing with conservation or epidemiological concerns, misidentified faecal samples from non-targeted species within the analysed sample can dramatically affect study conclusions. In the present study, 9% of the field-collected faecal samples were misidentified using morphological criterion. Hence, identifying the faeces origin with species-specific primer pairs and quantitative PCR^15,46^ is one alternative. However, the results presented here and elsewhere^26^ have shown that a positive detection in real-time PCR may reflect the contamination of samples with a small amount of DNA from targeted species. This can be illustrated through the human contamination usually detected in sequencing runs. In contrast, a positive species-specific PCR can partly reflect cross-contamination of samples or field contamination. Combining several replicates of PCR amplification with and without a blocking oligonucleotide would increase food item detection sensitivity per sample, particularly for small samples or when targeting the DNA of rare prey species. However, the number of primer mismatches and the concentration of blocking oligonucleotide can affect the detection of target species^35^.

In the present study, DNA metabarcoding allowed the identification of 90% of prey items up to species level and the remaining 10% to the genus level. It led to both a high detection rate and the high-resolution identification of closely related prey species (e.g. *M. arvalis* versus *M. agrestis*, *A. flavicollis* versus *A. sylvaticus*, Table S2). Using a DNA sequence database and bird-specific primer pairs to analyse faecal samples, Zarzoso-Lacoste and colleagues^47^ highlighted similar efficiency in using molecular ecology for the identification of bird species in cat and rat diets. Ordinarily, this level of precision is achieved only in cat diet studies that rely on prey animals brought back to the home (e.g. from 92% identified to at least class level^48^ to 50-90% identified to species level^49-51^), not with conventional methods analysing scat and stomach contents (e.g. from 25%^52^ to 59%^53^). Moreover, this molecular technic allows for the identification of a greater number of species than the conventional approach-based identification of prey items from scat and stomach contents used in previous cat diet studies (37 prey taxa in the present study versus from 3 to 14 different species^42,52-54^).

The results of this study demonstrate that DNA metabarcoding has the great advantage of allowing the detection of occurrences of food items that leave undetectable remains in the field (such as human-related food items) in a non-invasive sampling approach. This method gives insights on the potential origin of ingested food items (e.g., pet food or scavenged human-processed food versus scavenged carrion), with a taxonomic identification of 76% of vertebrate cyt*b* variants to the genus level for human-related food items (the remaining 24% to family, order or superorder levels). This makes this method useful for evaluating the exploitation of human-linked foods, and therefore it is possible to observe the full spectrum of cats’ diet, rather than concealing this food category due to inconvenience in detection^55^. So far, only Liberg^56^ has estimated the amount of household food in the diet of house cats as being the difference between the mass of meat required per day and a calculated amount of ingested prey. Nevertheless, the results of the calibration study make it clear that the rate of false negatives is partly linked to DNA integrity (and so to food item categories). This finding suggests that even if DNA metabarcoding allows more efficient detection of specific food items, certain items in the diet may remain highly underestimated (e.g. 46% for pet food and 69% for raw fish and meat), which could have a significant impact on conservation strategies. We encourage researchers to perform similar calibration studies and to be aware of these methodological challenges which could be species-specific. It should be noted, however, that sequence proportions matched biomass within a food category (e.g. rodents and birds).

To conclude, the present study joins several metabarcoding studies considering the biases and limitations of this approach, particularly in terms of using benchtop HTS platforms. Our findings indicate that the detection of degraded DNA in a heterogeneous mixture may vary accordingly to the degree of DNA degradation. This can drastically affect the identification of the ingested taxa for species having a diet possibly containing human-linked food items, and thus the inferences drawn from this diet, regarding the impacts of human activities on wildlife populations and food-web dynamics in anthropogenic landscapes. Nevertheless, performing a calibration study, generating sufficient numbers of technical replicates, using a sequencer such as MiSeq to produce a high quantity and quality of reads and applying appropriate metrics to summarise sequence data appears to be a robust approach for investigating the exploitation of a large range of food items for urban wildlife populations relying on a mixed-diet, above all in endangered species, for which it is preferable to use non-invasive survey approaches. Indeed, compared to conventional approaches based on morphological analysis, our DNA metabarcoding approach allows the identification of a large number of food items up to species level, including a wide range of food taxa leaving undetectable remains.

## Methods

### Faecal sample collection

#### Calibration study

A feeding trial was performed with a housebound cat fed a known diet during 22 consecutive days in May 2012 and 27 consecutive days in November 2012 (see Table S1 and Table S5 for details). The quantity of the daily diet of the housebound cat was constant but differed in composition from day to day. Daily rations included one or two of the following categories of food item: (1) processed wet or dry pet food (commercial meat-based cat food, Eco+ product line, SCAMARK, France); and (2a) prey items (immediately frozen after death) or (2b) surplus raw fish or meat products (supermarket withdrawn products with an early expiration date). The daily rations of individual prey species, processed pet food, and surplus raw fish and meat products were weighed and distributed evenly in three to four meals. We assumed that the probability of recovering DNA from these different food items depended on the type of ingested tissue (solid and soft tissues in whole prey carcasses versus only soft tissues in human-related food) and their quality (processed soft tissues in pet food versus raw soft tissues for surplus products). The faecal samples (*n* = 41) were then collected from a litter tray that was cleaned out daily, and these were put in labelled plastic bags and stored at −20 °C.

#### Field study

The faecal samples from free-ranging domestic cats were collected in a rural area located in north-eastern France (Ardennes region) as part of a joint study (Forin-Wiart et al. unpublished data). Cats’ faeces were sampled from a monitored free-ranging domestic cat population composed of 162 individuals, 40% of which were free-ranging outdoor cats fed *ad libitum* by their owners, and 60% of which were farm cats fed from time to time with milk, dry pet food or leftovers (see Forin-Wiart and colleagues^41^ for the census and cat feeding status definitions). A total of 359 fresh faecal samples were collected (1) along repeated transects within villages and isolated farms (total distance = 11 km) and linear transects crossing areas surrounding the villages (total distance = 15 km) and (2) from the litter trays of free-ranging individuals in June–July 2011 (*n* = 86 and *n* = 27, respectively), November– December 2011 (*n* = 101 and *n* = 25, respectively) and March 2012 (*n* = 108 and *n* = 12, respectively). Putative cat faecal samples were identified in the field by their shape and size^57^ prior to molecular identification based on cat-specific quantitative PCR (qPCR, see below). The 359 putative felid faecal samples were labelled with their location (using a GPS) and stored at −20 °C in plastic bags.

All experiments were performed in accordance with relevant guidelines and regulations.

### DNA extraction

All DNA extractions were performed at the SAS SPYGEN (France) laboratory in a room dedicated to degraded or rare DNA. Approximately 500 mg of faecal material was taken from all parts of the scat, as predator and prey DNA are not evenly distributed, then pooled, homogenised and 15 mg of this material was taken for DNA extraction, as recommended by Deagle and colleagues^26^. Total DNA was extracted following Shehzad and colleagues^58^. The QIAamp DNA Stool Mini Kit was used following the manufacturer’s instructions (Qiagen). The DNA extracts were recovered in a total volume of 250 µL. Ten extraction blanks (containing no faecal sample) were included as a negative control to test for contamination. DNA extraction success was evaluated by SAS SPYGEN with a cat species-specific designed primer (KiCqStart™ Primers) by following the universal SYBR Green quantitative real-time polymerase chain reaction PCR (qPCR) protocol (melt curve analysis of above amplification, melt peak = 81.5°C ± 0.5°C). This step was also used to validate the origin of field-collected faecal samples.

### DNA metabarcoding

A 136-bp mini-barcode of the cytochrome *b* gene was targeted, as it is effective in identifying a large range of vertebrates, including ungulates, rodents, birds and fish, from degraded samples^11,59,60^. We have verified *in silico* the absence of critical 3’-terminal mismatches between PCR primers and DNA targets using a sequence alignment of 1689 NCBI (National Centre for Biotechnology Information http://www.ncbi.nlm.nih.gov/) sequences from 44 species, corresponding to the species used in the calibration study and to rodent species from France (see Alignment S1 for the detailed list of species and Alignment file in fasta format).

Sequencing was successively performed using the Ion Personal Genome Machine (Ion PGM; Life Technologies) platform and the MiSeq (Illumina) platform to analyse the 400 DNA extractions obtained from the putative cat faecal samples (41 samples from the calibration study and 359 from the field study). Each DNA extraction was sequenced four times: twice independently on each HTS platform. A total of 5 negative extraction controls (in two replicates) were sequenced on both HTS platforms, as well as 2 and 9 negative PCR controls (in two replicates) by Ion PGM and MiSeq respectively. Due to the specificities of each platform, the DNA library preparation differed slightly (see below).

The Ion PGM libraries were prepared twice for each DNA extraction by amplifying the 136-bp fragment of cytochrome *b* described by Galan and colleagues^11^. We used the fusion primers L15411F (5’-CCA TCT CAT CCC TGC GTG TCT CCG ACT CAG NNNNNNN GAY AAA RTY CCV TTY CAY C-3’) and H15546R-PGM (5’CCT CTC TAT GGG CAG TCG GTG AT NNNNNNN AAR TAY CAY TCD GGY TTR AT-3’) modified in 5’ by the addition of individual-specific 7-bp MIDs (Multiplex IDentifiers NNNNNNN) and adaptors required for the emPCR and the Ion PGM sequencing. PCRs were performed following the procedure detailed by Galan and colleagues^11^. After PCR pooling and size selection by gel extraction, the amplicon libraries were sequenced by the company Genes Diffusion (Douai, France) on an Ion PGM system using an Ion 316 Chip Kit version 2 (Ion Torrent, Life Technologies).

The MiSeq libraries were prepared using a two-step PCR protocol (see Illumina Application Note Part #15044223) combined with the dual-index paired-end sequencing approach described by Kozich and colleagues^61^ due to the significant length of the Illumina adaptors (see details by Galan and colleagues^62^). During PCR1, the 136-bp fragment of cyt*b* was amplified twice for each DNA extraction with the primers modified in 5’ by the addition of a partial overhang Illumina adapter: L15411F-MiSeq 5’-TCG TCG GCA GCG TCA GAT GTG TAT AAG AGA CAG (none or C or TC or ATG) GAY A AA RTY CCV TTY CAY CC-3’ and H15546R-MiSeq 5’-GTC TCG TGG GCT CGG AGA TGT GTA TAA GAG ACA G (none or C or TG or GCT) AAR TAY CAY TCD GGY TTR AT-3’. The alternative bases between the partial adaptors and the target-specific primers corresponded to a 0-to 3-bp ‘heterogeneity spacer’ designed to mitigate the issues caused by low sequence diversity amplicons in Illumina sequencing^63^. These four versions of each primer were mixed before PCR1, which was performed in the same conditions as the Ion PGM PCR libraries, i.e. following the procedure detailed by Galan and colleagues^11^.

PCR1 was then used as the DNA template for PCR2, consisting of a limited-cycle amplification step to add multiplexing indices and Illumina sequencing adapters P5 and P7 at both ends of each DNA fragment. The primers P5 (5’-AAT GAT ACG GCG ACC ACC GAG ATC TAC AC NNNNNNNN TCG TCG GCA GCG TC-3’) and P7 (5’-CAA GCA GAA GAC GGC ATA CGA GAT NNNNNNNN GTC TCG TGG GCT CGG-3’) were synthesized with the different index sequences (NNNNNNNN) described by Kozich and colleagues^61^. PCR2 was carried out in a 10-µL reaction volume using 5 µL of PCR Multiplex Kit Master Mix (Qiagen) and 0.7 µM of each primer. One microliter of the PCR1 product was added to each well. PCR2 was started with an initial denaturation step of 95 °C for 15 min, followed by 5 cycles of denaturation at 94 °C for 40 s, annealing at 45 °C for 45 s and extension at 72 °C for 60 s, followed by a final extension step at 72 °C for 10 min. After PCR pooling and size selection by gel extraction, the amplicon library was sequenced at the local Plateau GPTR-GENOTYPAGE facility (Montpellier, France) on a MiSeq instrument by paired-end sequencing using a MiSeq 500 Cycle Kit version 2 (Illumina).

### Data processing

#### Bioinformatics and taxonomic identification

The paired-end reads 1 and 2 produced by MiSeq were assembled with the ‘make.contigs’ tool available in the mothur program^64^. This step enabled the removal of an important number of sequencing errors: the pairs of sequences were aligned and any positions with differences between the two reads were identified and corrected using the quality score of each read position. SESAME barcode software (SEquence Sorter & AMplicon Explorer^65^) was used for both Ion PGM and MiSeq data to sort the sequences and identify the variants at a taxonomic level (genus or species) using GenBank and three different similarity thresholds for BLAST+FASTA assignment (95%, 98% and 100%). The abundance table was then filtered.

#### Data filtering for false positive and negative occurrences using multiple controls

The arbitrary filtering procedure of high-throughput sequencing data to mitigate the effects of contamination on data analysis may lead to over-(loss of information, false negatives) or a lack of stringency (over interpretation, false positives)^27^. This is particularly true as features of each protocol (laboratory precautions, primer set efficiency, etc.) may influence the rate of bias. Hence, a fixed standardized-filtering threshold in metabarcoding process is misleading and this threshold should be protocol-dependent and based on internal controls and technical replicates. Here, one non-arbitrary threshold was applied for each sequencing run, to filter the false-positive assignments of reads (T_FA_) of a PCR product and a cross-validation procedure was used with technical replicates (see Galan and colleagues^62^ for more details).

The T_FA_ threshold was applied to filter the false-positive assignments of reads to a PCR product due to the generation of mixed clusters during the sequencing^66^. T_FA*i*_defines for each variant *i* the read number above which the PCR product is considered as a positive occurrence. This phenomenon was estimated in our experiment using the bank vole *Myodes glareolus* haplotype sequenced in parallel with the samples from the calibration study. As this rodent was handled separately from the faecal samples of the calibration study before sequencing, the presence of reads from *Myodes glareolus* in a sample of the calibration study indicated a sequence assignment error due to the Illumina sequencing (i.e., generation of mixed clusters). We determined the maximal number of reads of *Myodes glareolus* assigned to a PCR product. We then calculated the false-assignment rate (R_FA_) for this PCR product by dividing this number of reads by the total number of reads from *Myodes glareolus* in the sequencing run. Moreover, the number of reads for a specific variant *i* misassigned to a PCR product should increase with the total number of reads of this variant in the sequencing run. We therefore defined T_FA*i*_threshold as the total number of reads in the run for a variant *i* multiplied by R_FA_ (e.g. if an Ion PGM run produced 34,000 reads of an *Arvicola terrestris* haplotype and if R_FA_ = 0.05%, then T_FA*At*_= 17 reads for this *Arvicola terrestris* haplotype at the 100% similarity threshold). PCR products with less than T_FA*i*_for a particular variant *i* were considered negative. Lastly, we discarded not-replicated positive results for at least 50% of the PCR replicates to remove inconsistent variants due to PCR or sequencing errors or unconfident variants that could be associated with the remaining false positive results^28^. Following these data filtering stages, a table of occurrences of observed food items per sequencing run was obtained in order to compare the performance of the two platforms (see below). In a final step, the composition of each faecal sample and the food occurrence data from both HTS platforms was pooled to summarise the effectiveness of this DNA metabarcoding approach from a dietary study perspective.

### Data analysis

Model selection was based on the Information Theoretic approach and was performed with the Akaike Information Criterion (AIC) following the model selection approach described by Zuur and colleagues^67^. All statistical analyses were conducted with the statistical software R 3.3 (R Development Core Team 2016^68^), and a value of alpha = 0.05 was used for all tests.

#### Calibration study

The ability of the two HTS platforms to detect consumed items from a diet of no processed food (prey items only) or from different degrees of processed food (wet and dry pet food, raw fish or meat) was assessed by calculating the frequency of occurrence metric. For each HTS platform and each food item category (prey, pet food or raw fish and meat), the number of faecal samples containing haplotype sequences that were informative enough to assign a food item (observed occurrence) was compared to the minimum expected occurrence, in order to identify the sensitivity and bias of each sequencing strategy. The minimum expected occurrence was determined by considering that the DNA associated with a given food item ingested by the housebound cat was expected to be detected in the scat emitted at least 12 h after the ingestion of the food item, according to the transit time previously assessed by Forin-Wiart and colleagues^41^. The agreement of food item detection (i.e. its occurrence in one, both or none of the replicates from a sequencing run) between each HTS platform and each successfully amplified scat was also estimated. Additionally, for mixed meals (i.e. including two food item categories), the proportion of each food item category by mass ingested by the housebound cat, relative to the total mass of the food ingested, was compared to the proportion of each food item category identified by haplotype information in the sequence reads, relative to the total number of reads of a specific faecal sample. This measure allowed the investigation of whether the proportion of a given food item category in the diet of cats was reflected in the HTS sequence counts. These proportions were compared using Fisher’s exact tests for count data.

#### Field study

Small- and medium-sized rodent, insectivore, wild bird and reptile DNA in field-collected samples were placed in the ‘Prey items’ category and its sub-categories (see Table S2 for details). In contrast, DNA from domestic ungulates (cow, goat, sheep, pig) and farmed birds (chicken, duck, guinea fowl, turkey), as well as farmed fish and game species (*Capreolus capreolus, Cervus elaphus,* Perdicinae *spp*, and rabbit *Oryctolagus cuniculus*) were placed in the ‘Human-linked food’ category and its sub-categories (see Table S2 for details). This latter category was assumed to originate from pet food, leftovers or hunting scraps which do not contain preserved hard remains unlike prey items. Depending on the study area location, one food item could originate from one to three different sources (e.g. rabbits can be actively hunted by cats when available in the area, scavenged from human leftovers or found in pet food). During our experiment, we considered that game species were only available from pet food or human leftovers (rabbit hutches and hunting scraps) for cats in this area. These food item categories were defined for ecological (understand the relative contributions of wildlife versus food items provided by or scavenged from humans for further dietary studies) and technical reasons (fit with the composition of food items (whole carcass versus processed food versus raw items) which affects the detectability of DNA).

One or more than one anthropogenic food taxa can be identified in one faecal sample, i.e. multiple occurrences, without being able to associate taxa to independent ingestion events (pet food is composed of a mix of meat and/or fish belonging to various taxa^69^) or conversely to prey taxa and game species occurrences. Accordingly, for comparison between food categories or sub-categories, occurrence data, i.e. presence/absence in one or two replicates per HTS platform, were used (e.g. If one faecal sample was composed of two anthropogenic food taxa, including one taxa in two replicates and the other one in one replicate and two preys taxa, both in replicates. Then one observed occurrence in two replicates will be given to the anthropogenic food category, as well as to the prey items category).

To identify the sensitivity of each sequencing protocol, the probability of detecting a food item in field-collected samples, i.e. presence/absence (Table S3), was modelled as a function of the HTS platform (fPlatform), the food item category (fFoodItem, reptile occurrence was excluded for modelling purposes) and the interaction between fPlatform and fFoodItem, using a Generalized Linear Model (GLM) with a binomial link function. The fit of the full model (lme4 package^70707070^) and the reduced model was compared using a deviance test (G²) as an alternative to an individual parameter-based approach by considering fSample as a random effect. Multiple comparisons with interaction terms were then adjusted using Tukey’s post-hoc analyses (lsmeans package^71717171^).

Finally, multiple occurrences for prey and human-linked food taxa, respectively were conserved to summarise the effectiveness of our DNA metabarcoding approach for further dietary study perspectives by calculating the frequency of occurrence metric.

## Acknowledgements

MAFW was supported by a PhD grant (Région Champagne-Ardenne and Conseil Général des Ardennes). We are grateful to the people in the study area who allowed us to prospect for and collect samples, to the cat owners who collected samples for the study, to Matthieu Bastien, Ivan Puga Gonzales and Nancy Rebout for their statistical advice, to Vincent Viblanc for its constructive comments, and to Peter Stuart for editing the manuscript. We also sincerely thank Spécialités Pet Food (SPF) for funding the DNA extraction and the two HTS runs.

## Author Contributions

MAFW and MLP contributed to the study design and sample collection. MG conducted the molecular analysis. MAFW, MG and SP conducted the bioinformatics analysis. MAFW conducted the data analysis, performed the statistical analysis and prepared the tables and graphs. MAFW, MLP and MG led the writing of the manuscript. All authors contributed critically to the drafts and gave final approval for publication.

## Additional Information

The authors declare no competing interests.

## Data availability

Should the manuscript be accepted, the data supporting the results (raw sequence reads (fastq format), sequence alignment (fastq format), raw table of abundance and filtered table of occurrence including taxonomic affiliations) will be published along with the manuscript in a Dryad Digital Repository.

## References

1 McCleery, R. A., Moorman, C. E. & Peterson, M. N. Urban wildlife conservation: theory and practice. (Springer, 2014).

2 Murray, M. et al. Greater consumption of protein□poor anthropogenic food by urban relative to rural coyotes increases diet breadth and potential for human-wildlife conflict. Ecography 38, 1235–1242 (2015).

3 Robardet, E. et al. Infection of foxes by Echinococcocus multilocularis in urban and suburban areas of Nancy, France: influence of feeding habits and environment. Parasite 15, 77–85 (2008).

4 Newsome, S. D., Ralls, K., Job, C. V. H., Fogel, M. L. & Cypher, B. L. Stable isotopes evaluate exploitation of anthropogenic foods by the endangered San Joaquin kit fox (Vulpes macrotis mutica). J. Mammal. 91, 1313–1321, doi:10.1644/09-mamm-a-362.1 (2010).

5 Bateman, P. W., Fleming, P. A. & Le Comber, S. Big city life: carnivores in urban environments. J. Zool. 287, 1–23, doi:10.1111/j.1469-7998.2011.00887.x (2012).

6 Pompanon, F. et al. Who is eating what: Diet assessment using next generation sequencing. Mol. Ecol. 21, 1931–1950 (2012).

7 Yoccoz, N. G. The future of environmental DNA in ecology. Mol. Ecol. 21, 2031–2038 (2012).

8 Egeter, B., Bishop, P. J. & Robertson, B. C. Detecting frogs as prey in the diets of introduced mammals: a comparison between morphological and DNA-based diet analyses. Molecular Ecology Resources 15, 306–316, doi:10.1111/1755-0998.12309 (2015).

9 Taberlet, P., Coissac, E., Pompanon, F., Brochmann, C. & Willerslev, E. Towards next-generation biodiversity assessment using DNA metabarcoding. Mol. Ecol. 21, 2045–2050 (2012).

10 Shokralla, S., Spall, J. L., Gibson, J. F. & Hajibabaei, M. Next generation sequencing technologies for environmental DNA research. Mol. Ecol. 21, 1794–1805 (2012).

11 Galan, M., Pagès, M. & Cosson, J.-F. Next-generation sequencing for rodent barcoding: species identification from fresh, degraded and environmental samples. PLoS ONE 7, e48374, doi:10.1371/journal.pone.0048374 (2012).

12 Valentini, A., Pompanon, F. & Taberlet, P. DNA barcoding for ecologists. Trends Ecol. Evol. 24, 110–117, doi:10.1016/j.tree.2008.09.011 (2009).

13 Coghlan, M. L. et al. Metabarcoding avian diets at airports: Implications for birdstrike hazard management planning. Investigative Genetics 4 (2013).

14 De Barba, M. et al. DNA metabarcoding multiplexing and validation of data accuracy for diet assessment: Application to omnivorous diet. Molecular Ecology Resources 14, 306–323 (2014).

15 Shehzad, W. et al. Carnivore diet analysis based on next-generation sequencing: Application to the leopard cat (Prionailurus bengalensis) in Pakistan. Mol. Ecol. 21, 1951–1965 (2012).

16 Brown, D. S., Ebenezer, K. L. & Symondson, W. O. C. Molecular analysis of the diets of snakes: Changes in prey exploitation during development of the rare smooth snake Coronella austriaca. Mol. Ecol. 23, 3734–3743 (2014).

17 Brown, D. S., Jarman, S. N. & Symondson, W. O. C. Pyrosequencing of prey DNA in reptile faeces: Analysis of earthworm consumption by slow worms. Molecular Ecology Resources 12, 259–266 (2012).

18 Galal-Khallaf, A., Osman, A. G., Carleos, C. E., Garcia-Vazquez, E. & Borrell, Y. J. A case study for assessing fish traceability in Egyptian aquafeed formulations using pyrosequencing and metabarcoding. Fisheries Research 174, 143–150 (2016).

19 Galan, M. et al. 16S rRNA amplicon sequencing for epidemiological surveys of bacteria in wildlife: the importance of cleaning post-sequencing data before estimating positivity, prevalence and co-infection. mSystems 1, e00032–00016, doi:doi:10.1128/mSystems.00032-16 (2016).

20 Ficetola, G. F. et al. Replication levels, false presences and the estimation of the presence/absence from eDNA metabarcoding data. Molecular Ecology Resources 15, 543–556 (2015).

21 D’Amore, R. et al. A comprehensive benchmarking study of protocols and sequencing platforms for 16S rRNA community profiling. BMC Genomics 17, 55 (2016).

22 Sharma, N., Thind, S. & Sharma, D. Effect of meat processing on genomic DNA quality and specific gene amplification. Journal of Applied Animal Research 28, 69–72 (2005).

23 Coissac, E., Riaz, T. & Puillandre, N. Bioinformatic challenges for DNA metabarcoding of plants and animals. Mol. Ecol. 21, 1834–1847, doi:10.1111/j.1365-294X.2012.05550.x (2012).

24 Fitzgerald, B. M. & Turner, D. C. in The domestic cats: The Biology of its Behaviour (eds D.C. Turner & P. Bateson) 151–175 (Cambridge Univ. Press, 2000).

25 Spotte, S. in Free-ranging cats: Behavior, ecology, management (ed Wiley Blackwell) Ch. 9, 181–213 (John Wiley & Sons, 2014).

26 Deagle, B. E., Kirkwood, R. & Jarman, S. N. Analysis of Australian fur seal diet by pyrosequencing prey DNA in faeces. Mol. Ecol. 18, 2022–2038, doi:10.1111/j.1365-294X.2009.04158.x (2009).

27 Corse, E. et al. A from benchtop to desktop workflow for validating HTS data and for taxonomic identification in diet metabarcoding studies. Molecular Ecology Resources 17, e146–e159 (2017).

28 Ficetola, G. F., Taberlet, P. & Coissac, E. How to limit false positives in environmental DNA and metabarcoding? Molecular Ecology Resources 16, 604–607 (2016).

29 De Battisti, C. et al. Pyrosequencing as a tool for rapid fish species identification and commercial fraud detection. J. Agric. Food Chem. 62, 198–205 (2013).

30 Loman, N. J. et al. Performance comparison of benchtop high-throughput sequencing platforms. Nature Biotechnology 30, 434–439 (2012).

31 Deagle, B. E., Thomas, A. C., Shaffer, A. K., Trites, A. W. & Jarman, S. N. Quantifying sequence proportions in a DNA-based diet study using Ion Torrent amplicon sequencing: Which counts count? Molecular Ecology Resources 13, 620–633 (2013).

32 Egge, E. et al. 454 pyrosequencing to describe microbial eukaryotic community composition, diversity and relative abundance: a test for marine haptophytes. PloS one 8, e74371 (2013).

33 Pochon, X., Bott, N. J., Smith, K. F. & Wood, S. A. Evaluating detection limits of next-generation sequencing for the surveillance and monitoring of international marine pests. PLoS One 8, e73935 (2013).

34 Kartzinel, T. R. et al. DNA metabarcoding illuminates dietary niche partitioning by African large herbivores. Proceedings of the National Academy of Sciences of the United States of America 112, 8019–8024, doi:10.1073/pnas.1503283112 (2015).

35 Piñol, J., Mir, G., Gomez-Polo, P. & Agustí, N. Universal and blocking primer mismatches limit the use of high-throughput DNA sequencing for the quantitative metabarcoding of arthropods. Molecular Ecology Resources 15, 819–830, doi:10.1111/1755-0998.12355 (2015).

36 Deagle, B. E., Chiaradia, A., McInnes, J. & Jarman, S. N. Pyrosequencing faecal DNA to determine diet of little penguins: is what goes in what comes out? Conserv. Genet. 11, 2039–2048 (2010).

37 Deagle, B. et al. Counting with DNA in metabarcoding studies: how should we convert sequence reads to dietary data? bioRxiv, doi:10.1101/303461 (2018).

38 Deagle, B. E., Jarman, S. N., Coissac, E., Pompanon, F. & Taberlet, P. DNA metabarcoding and the cytochrome c oxidase subunit I marker: not a perfect match. Biol. Lett. 10, 20140562 (2014).

39 Sheppard, S. K. et al. Detection of secondary predation by PCR analyses of the gut contents of invertebrate generalist predators. Mol. Ecol. 14, 4461–4468 (2005).

40 Antton, A., Ostaizka, A., P., G. M. T. & Kristine, B. Scrutinizing key steps for reliable metabarcoding of environmental samples. Methods in Ecology and Evolution 9, 134–147, doi:doi:10.1111/2041-210X.12849 (2018).

41 Forin-Wiart, M.-A., Gotteland, C., Gilot-Fromont, E. & Poulle, M.-L. Assessing the homogeneity of individual scat detection probability using the bait-marking method on a monitored free-ranging carnivore population. European Journal of Wildlife Research 60, 665–672, doi:10.1007/s10344-014-0833-0 (2014).

42 Krauze-Gryz, D., Gryz, J. & Goszczyński, J. Predation by domestic cats in rural areas of central Poland: an assessment based on two methods. J. Zool. 288, 260–266, doi:10.1111/j.1469-7998.2012.00950.x (2012).

43 Pires, M. M., Widmer, C. E., Silva, C. & Setz, E. Z. F. Differential detectability of rodents and birds in scats of ocelots, Leopardus pardalis (Mammalia: Felidae). Zoologia (Curitiba) 28, 280–283 (2011).

44 Deagle, B. et al. Molecular scatology as a tool to study diet: analysis of prey DNA in scats from captive Steller sea lions. Mol. Ecol. 14, 1831–1842 (2005).

45 Elbrecht, V. & Leese, F. PrimerMiner: an r package for development and in silico validation of DNA metabarcoding primers. Methods in Ecology and Evolution 8, 622–626, doi:doi:10.1111/2041-210X.12687 (2017).

46 Knapp, J., Umhang, G., Poulle, M.-L. & Millon, L. Development of a real-time PCR for a sensitive one-step coprodiagnosis allowing both the identification of carnivore feces and the detection of Toxocara spp. and Echinococcus multilocularis. Applied and environmental microbiology 82, 2950–2958 (2016).

47 Zarzoso-Lacoste, D. et al. Improving morphological diet studies with molecular ecology: An application for invasive mammal predation on island birds. Biol. Conserv. 193, 134–142, doi:10.1016/j.biocon.2015.11.018 (2016).

48 Kauhala, K., Talvitie, K. & Vuorisalo, T. Free-ranging house cats in urban and rural areas in the north: useful rodent killers or harmful bird predators? Folia Zoologica 64 (2015).

49 Baker, P. J., Molony, S. E., Stone, E., Cuthill, I. C. & Harris, S. Cat about town: is predation by free-ranging pet cats Felis catus likely to affect urban bird populations? Ibis 150, 86–99 (2008).

50 Tschanz, B., Hegglin, D., Gloor, S. & Bontadina, F. Hunters and non-hunters: skewed predation rate by domestic cats in a rural village. European Journal of Wildlife Research 57, 597–602 (2011).

51 Krauze-Gryz, D., Żmihorski, M. & Gryz, J. Annual variation in prey composition of domestic cats in rural and urban environment. Urban Ecosystems 20, 945–952, doi:10.1007/s11252-016-0634-1 (2017).

52 Germain, E., Ruette, S. & Poulle, M.-L. Likeness between the food habits of European wildcats, domestic cats and their hybrids in France. Mammalian Biology 74, 412–417 (2009).

53 Biró, Z., Lanszki, J., Szemethy, L., Heltai, M. & Randi, E. Feeding habits of feral domestic cats (Felis catus), wild cats (Felis silvestris) and their hybrids: trophic niche overlap among cat groups in Hungary. Journal of Zoology (London) 266, 187–196 (2005).

54 Weber, J.-M. & Dailly, L. Food habits and ranging behaviour of a group of farm cats (Felis catus) in a Swiss mountainous area. Journal of Zoology (London) 245, 234–237 (1998).

55 Turner, D. C. in The domestic cat: the biology of its behaviour (eds D.C. Turner & P. Bateson) 63–70 (Cambridge University Press, 2014).

56 Liberg, O. Food habits and prey impact by feral and house-based domestic cats in a rural area in southern sweden. J. Mammal. 65, 424–432 (1984).

57 Corbett, L. K. Feeding ecology and seasonal organization of wildcats (Felis silvestris) and domestic cats (Felis catus) in Scotland., PhD thesis, University of Aberdeen, Scotland, (1979).

58 Shehzad, W. et al. Prey Preference of Snow Leopard (Panthera uncia) in South Gobi, Mongolia. PlOS ONE 7, e32104, doi:10.1371/journal.pone.0032104 (2012).

59 Kappel, K., Haase, I., Käppel, C., Sotelo, C. G. & Schröder, U. Species identification in mixed tuna samples with next-generation sequencing targeting two short cytochrome b gene fragments. Food Chem. 234, 212–219 (2017).

60 Teletchea, F., Bernillon, J., Duffraisse, M., Laudet, V. & Hänni, C. Molecular identification of vertebrate species by oligonucleotide microarray in food and forensic samples. J. Appl. Ecol. 45, 967–975 (2008).

61 Kozich, J. J., Westcott, S. L., Baxter, N. T., Highlander, S. K. & Schloss, P. D. Development of a dual-index sequencing strategy and curation pipeline for analyzing amplicon sequence data on the MiSeq Illumina sequencing platform. Appl. Environ. Microbiol. 79, 5112–5120 (2013).

62 Galan, M. et al. Metabarcoding for the parallel identification of several hundred predators and their prey: Application to bat species diet analysis. Molecular Ecology Resources 18, 474–489, doi:10.1111/1755-0998.12749 (2018).

63 Fadrosh, D. et al. An improved dual-indexing approach for multiplexed 16S rRNA gene sequencing on the Illumina MiSeq platform. Microbiome 2, 6 (2014).

64 Schloss, P. D. et al. Introducing mothur: Open-Source, Platform-Independent, Community-Supported Software for Describing and Comparing Microbial Communities. Appl. Environ. Microbiol. 75, 7537–7541, doi:10.1128/aem.01541-09 (2009).

65 Piry, S., Guivier, E., Realini, A. & Martin, J. F. | SE| S| AM| E| Barcode: NGS-oriented software for amplicon characterization–application to species and environmental barcoding. Molecular Ecology Resources 12, 1151–1157 (2012).

66 Kircher, M., Sawyer, S. & Meyer, M. Double indexing overcomes inaccuracies in multiplex sequencing on the Illumina platform. Nucleic Acids Res. 40, e3–e3, doi:10.1093/nar/gkr771 (2012).

67 Zuur, A., Ieno, E. N., Walker, N., Saveliev, A. A. & Smith, G. M. Mixed effects models and extensions in ecology with R. (Springer, 2009).

68 R: A language and environment for statistical computing. (Vienna, Austria, 2016).

69 Okuma, T. A. & Hellberg, R. S. Identification of meat species in pet foods using a real-time polymerase chain reaction (PCR) assay. Food Control 50, 9–17, doi:https://doi.org/10.1016/j.foodcont.2014.08.017 (2015).

70 Bates, D., Mächler, M., Bolker, B. & Walker, S. Fitting linear mixed-effects models using lme4. arXiv preprint arXiv:1406.5823 (2014).

71 Lenth, R. V. Least-Squares Means: The R Package lsmeans. Journal of Statistical Software 69, 1–33, doi:10.18637/jss.v069.i01 (2016).

